# Recent bloom of filamentous algae in Lake Baikal is caused by *Spirogyra* Link., 1820 of local origin

**DOI:** 10.1101/2020.02.10.942979

**Authors:** Elena Mincheva, Tatiana Peretolchina, Tatyana Triboy, Yrij Bukin, Luybov Kravtsova, Andrey Fedotov, Dmitry Sherbakov

## Abstract

Molecular phylogeny inferred from *rbc*L nucleotide sequences obtained from the single sterile filaments of green algae collected around the perimeter of Lake Baikal indicates the polyphyletic origin of the representatives of genus *Spirogyra* Link., 1820 inhabiting the lake. The common ancestor of all Baikal Spirogyra dates back at least to 20 MYA. This roughly coincides with the age of continuously existing freshwater body in the confines of current Baikal. The descendants of this node include both Baikal and non-Baikal species and thus suggesting a complex history of multiple emigrations and immigrations. There is at least one major lineage of the Baikal *Spirogyra* in the phylogeny descending uninterruptedly from the common ancestor of all *Spirogyra* species found so far in the lake. The likely explanation is its permanent presence in the ecosystem. All this allows us to hypothesize that the current bloom is a spectacular but natural response of the Baikal ecosystem to the increased pollution.

## Introduction

High production of filamentous algae is among the most obvious and dramatic manifestations of unwanted changes in different kinds of aquatic ecosystems [1–3]. It may take place both in marine and freshwater environments often in response to an elevated influx of nutrients or other impacts promoting production [4]. Although the cases of economic exploitation of filamentous algae are known [5], usually their mass occurrence in an ecosystem is regarded as the indicator of ecological damage going on [6–9]. Recently the usual temporal and spatial patterns of green algal growth were spoiled dramatically in Lake Baikal (East Siberia) [10–14]. Since Lake Baikal is the most ancient and the largest of giant freshwater lakes [15] containing ca. 20% of the world’s fresh water and home for highly diverse and endemic biota, this development has caused strong public and scientific concern. However, the very scale of Lake Baikal and the diversity of its biotopes offer an additional challenge to the study of its ecosystem changes. Indeed, different regions of the lake range from highly polluted spots permanently suffering under strong human impact to pristine areas where no ecological changes still may be detected [11, 12]. Therefore the results obtained for one locality are hardly expandable to the other parts of the same lake. Previous studies had demonstrated that several of filamentous algal species belonging to genus *Spirogyra* comprise large if not a major part of the algal blooms in the polluted parts of Lake Baikal [13]. Several species of *Spirogyra* were found in shallow bays of Lake Baikal as rare filaments [16]. This makes it possible to hypothesize, that the blooming *Spirogyra* could originate from those few surviving opportunists exploiting an increased influx of nutrients. The alternative hypothesis proposes that the blooms were caused by exotic invaders using the window of opportunity existing due to anthropogenic impact. We believe that the expected further development of ecological transformations of the Baikal ecosystem depends on which hypothesis is true with a much more catastrophic perspective in the latter case.

Since the first studies revealed that the algal community of the green sludge consists of many different filamentous algae, we have adopted the approach based on the identification of single filaments. Therefore in order to elucidate the evolutionary roots of *Spirogyra* we have isolated individual algal filaments from 18 localities scattered along the shore of the lake. The barcode chloroplast gene *rbc*L was used to provide phylogenetic evidence for Baikal origin of the most abundant species of *Spirogyra* and to demonstrate the existence of complicated immigration/emigration patterns of its relation with non-Baikal habitats.

## Materials and methods

Samples of filamentous Charophyta were collected by divers at the 18 localities along the perimeter of Lake Baikal during the three expeditions of RV “Titov” from 2016 to 2019 (Fig.1, Table S1).

**Fig. 1.**
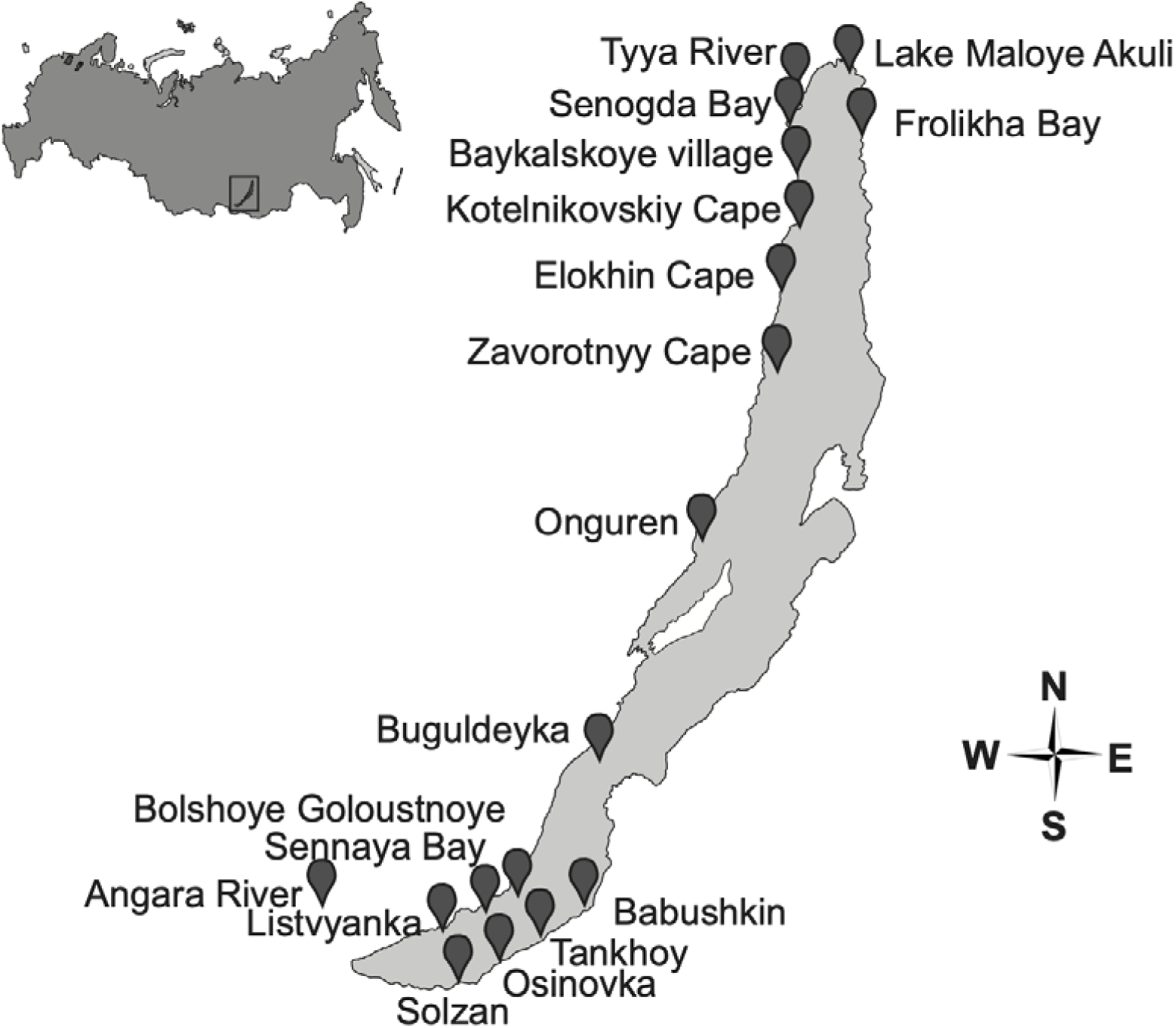
Map of localities where filamentous green algae were sampled.

Separate sterile (no conjugation) filaments of *Spirogyra* were manually picked up under a microscope and placed in tubes with 2% CTAB and frozen at−20*°*C.

DNA was extracted according to the modified protocol by Doyle and Dickson [17].

Fragments of *rbc*L were amplified using primers rbcL183 M28F 5’-GGTGTTGGATTTAAAGCTGGTGT-3’ and *rbc*L M1390R 5’-CTTTCAAAYTTCACAAGCAGCAG-3’ [18]. 35 cycles were preceded with 4 min. long pre-denaturation at 95°C and were 20 sec. at 95*°*C, 20 sec. min at 55*°*C, 1.5 min 72*°*C (5 min at the last cycle). For the second stage, we have developed the *Spirogyra*–specific primers Spir_rbcL_1340R 5’-CTAACTCAGGACTCCATTTG-3’, Spir_rbcL_330F 5’-CTATTGTAGGTAACGTATTTGG-3’, which were used with the same protocol as at stage 1. These primers flank the most variable part of *rbc*L about 1kB long. PCR amplification of the respective fragments was performed with the Biolabmix (Novosibirsk, Russia) HS180Taq kit in 25 *μ*l of the reaction mixture in a Bio-Rad (USA) thermocycler Bio-Rad T100. After electrophoresis purification and extraction with the gel elution kit (Biosilica Co., Novosibirsk, Russia) the direct sequencing of forward and reverse sequences were performed in Research and Production Company SYNTOL (Russia) using an ABI 3130 automated sequencer.

### Phylogenetic inferences

DNA sequences of *rbc*L obtained in this study (Table 1 in Supplemantary), were merged together with the sequences of *Spirogyra* available in NCBI database [19-21], the alignment was checked with mafft [22, 23]. Three species of other Charales (*Zygogonium ericetorium, Zygnema cylindricum*, and *Mougeoutia* sp.) were used as the outgroup rooting the phylogeny. The phylogenetic inferences were performed with IQ-TREE [24] with optimal model of molecular evolution (TIM3+I+G4) chosen according to its BIC value, the robustness of the topology was tested both using ultra-fast bootstrap [25] and SH-alrt test [26].

BEAST ver.1.8.4 [27] was used for Bayesian estimations of the ages of nodes numbered on Fig.2a. The ML-tree inferred with IQ-TREE was used as the user-tree, as well as the TIM3+I+G4 model chosen by IQ-TREE was assumed *a priori*. The logarithmically relaxed model of the molecular clock [28] was used together with the evolutionary birth-death model [29]. As the source of calibration, the monophyletic clades of Hawaiian *Spirogyra* restricted to single islands were used under the assumption of approximately simultaneous appearance of the island and expansion of *Spirogyra* to its water bodies [30]. Similar dating strategy was successfully used in [31, 32].

The same tree was used for studies of evolutionary parameters of the distribution of the *Spirogyra* unique haplotypes Mesquite v.3.6 [33].

**Fig. 2.**
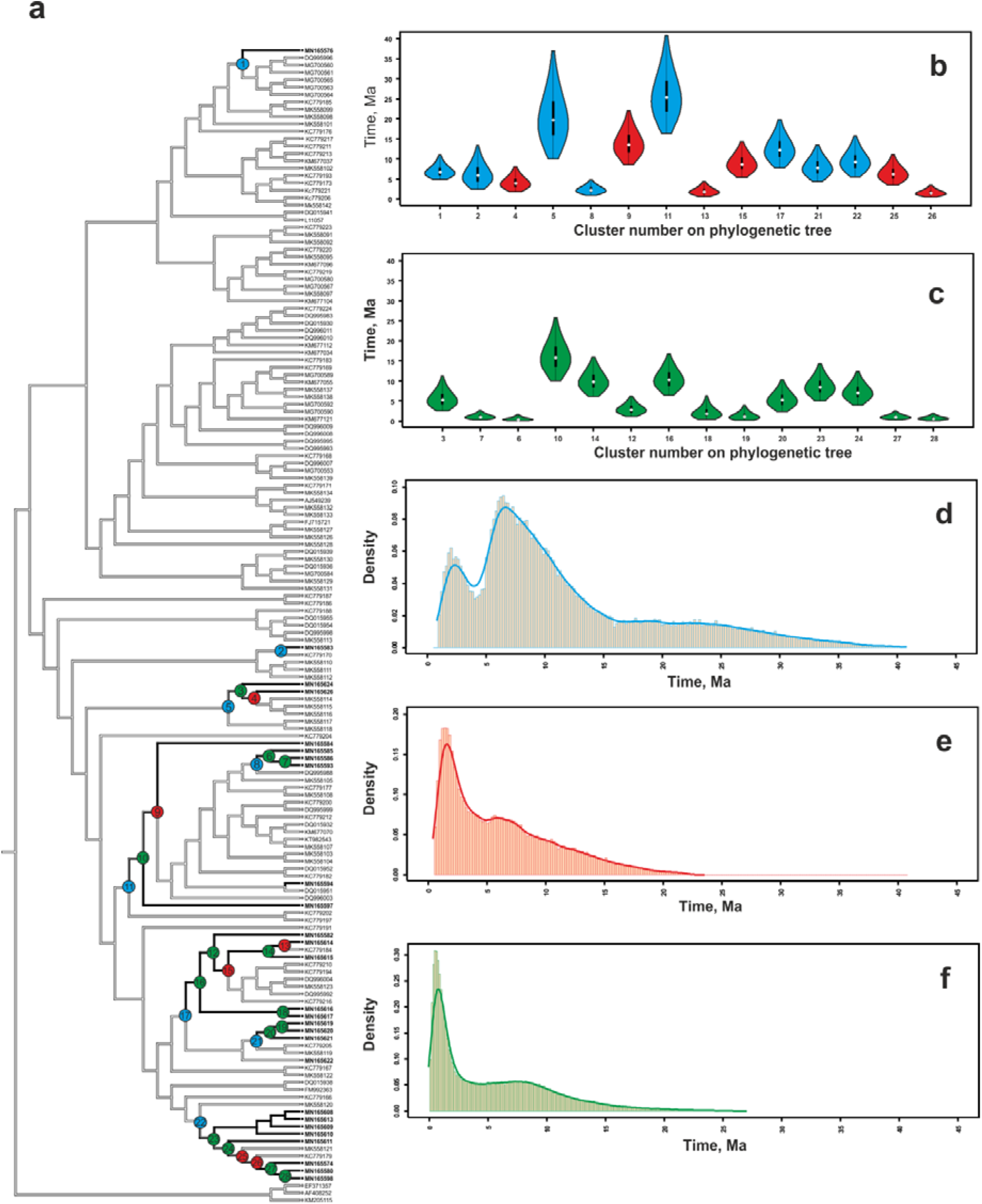
Evolution of the distribution of Spirogyra between Lake Baikal and other water bodies. A. Ultrametric BEAST tree estimated from the maximum likelihood tree inferred from *rbc*L sequences (Table S1 and Fig.S1). The blue square outlines the major lineage containing all but one haplotypes of the Baikal samples. Numbered circles denote tree nodes for which time estimates have been performed. Blue circles indicate immigration events, red circles indicate emigrations, and the green ones indicate nodes with all descendants remaining in Baikal. B. and C. Time estimates for the nodes outlined at panel A. Violin plots cover 95% confidence limits. D, E, and F: Probability distributions for D) no migration nodes, E) emigration from Baikal; F) immigration into Baikal

### Phylogeographic analysis

The phylogeographic analysis was based on all sequence data available on *Spirogyra* by the fall of 2019. Unfortunately, the sampling localities are known for less than a half of *rbc*L sequences published. Therefore we have approximated the geographic distribution by a binary trait “Baikal” vs. “non-Baikal” and used both parsimony and likelihood approaches to describe the present and past trends in the evolution of this trait.

The amount of long-term isolation between Baikal and not Baikal haplotypes on an evolutionary time-scale was estimated by permutation test on the tree topology inferred from nucleotide sequences [33]. The tree was subjected to 150 random branch-and-bounds, after which different metrics of the amount of phylogenetic signals such as retention index or the parsimonious number of steps. This procedure was repeated for 10^6^ times to compromise between the vast size of tree space and the probability of repeated samplings.

## Results and discussion

The main focus of the study was at the origin of the mass haplotypes present in Lake Baikal: are they the recent invaders having their roots outside the lake, or they are the endemic opportunists originating from previously rare species found there in the middle of XX century [16].

Phylogenetic inference performed for 366 *rbc*L sequences (157 unique haplotypes) resulted in a fully resolved tree with highly supported most if internal nodes (Fig.2).

The most important feature of the tree is that it splits into two major lineages, one of which contains the deep node ancestral to all Baikal sequences but a single one. The descendants of this node contain both Baikal OTUs and OTUs of American, Japanese, Hawaiian and European origin.

Up to date, *Spirogyra* diversity is studied in four localities: California [19], Hawaii [20], Japan [21] and Lake Baikal (current study). Other sequences clustering within the genus and presented in the NCBI database were treated as being of unknown origin. The ML phylogeny inferred (Fig. S1) suggests that each of the regions was populated by *Spirogyra* species more than once. All possible directions of past migration were found except for the direct migration between Lake Baikal and Hawaii.

Likelihood ratio test for the hypothesis of asymmetry vs. symmetry of transition between “Baikal” - “non-Baikal” treated as a binary character gives 6.98454086. Assuming *χ*^2^ df = 1, this favors the «asymmetric» hypothesis with rejection level at *p* = 0.0082. It was found, that the exchange with the outside world was biased (bias ≈ 0.13833) towards immigration to Lake Baikal. Never the less, emigration must be significant since none of the two unidirectional models cannot explain the current distribution of *Spyrogira* if the phylogeny is accepted. Note that due to the high dispersal potential [34] the distribution of *Spyrogira* species is likely to be at equilibrium at any time. However, only one haplotype was found both in Lake Baikal and other fresh water bodies. This indicates noticeable difficulties in long-term successful immigration to Lake Baikal of the new *Spirogyra*. This finding contradicts strongly the common opinion of recent invasion to Lake Baikal by some exotic representatives of Charophyta.

The results of permutation test (Fig. S2) suggest a significantly high amount of phylogeny signal, making the change of state in the distribution character less probable than speciation.

The Bayesian approach and based on geological dating of the appearance times of Hawaiian Islands allowed to estimate the most likely substitution rate for the *rbc*L to 0.011% for million years with 95% confidence limits from 0.0063% to 0.016% per million years. The common ancestor of “Baikal” sequences dates back approximately to 18MYA which approaches ca. 20 – 25MYA which are regarded as the age of large freshwater lake permanently existing in confines of the current lake [15]. After that, approximately the first half of the history of the lake was dominated by the immigration of different lineages of *Spirogyra* followed by a long period of both emigrations and immigrations. Interestingly, the period preceding the cooling down of climate was very favorable for *Spirogyra* judging by the increase of the intralacustrine divergence (Fig. 2f) followed by a decrease in Pleistocene. This corroborates with the data by Izhboldina and coauthors [16, 35] who found the representatives of this genus as very rare single filaments in shallow waters of Lake Baikal in the middle of XX century. Thus the recent blooms appear to be a reverse of the long trend.

The results reported here indicate that it is likely that 1) multiple *Spirogyra* species were present in confines of current Lake Baikal since the appearance of a large fresh continuously existing water body ca. 25MYA. 2)There were multiple immigrations and emigrations from the outside, but establishing permanent lineages inside the lake was relatively rare making the Baikal algae noticeably isolated; 3)The *Spirogyra* species blooming recently in Lake Baikal are most likely local opportunists rather than successful exots exploiting an increase in nutrient influxes. We believe that taking into account long persistence and high variation in numbers over a short period of time, our findings presented here, allow us to propose, that the blooms of filamentous green algae may at least partially explain why relatively long “diatomless” periods in the well-studied diatom record of paleo-climate of Lake Baikal [36, 37] did not leave noticeable traces in evolutionary histories of Baikal benthic animals [38, 39].

