## Supplementary material for "Recent bloom of filamentous algae in Lake Baikal is caused by *Spirogyra* Link., 1820 of local origin": Supplemantal Table and Figures

**Table S1.** List of localities of *Spirogyra* samples were collected

| Strain | Locality and habitat, data of collection | New *rbc*L genes used for present phylogenetic analysis |
|---|---|---|
| Senogda47 | Lake Baikal, Senogda Bay (55.56 N 109.23 E) (17 Aug 2016) | MN165574 |
| Zavorotnyiy_44 | Lake Baikal, Zavorotnyy Cape (54.28 N 108.49 E) (15 Aug 2016) | MN165575 |
| Babuskin_46c | Lake Baikal, Babushkin (51.72 N 105.85 E) (29 Sep 2017) | MN165576 |
| Sludyanka_47g | Lake Slyudyanskoye (55.46 N 109.16 E) (16 Aug 2016) | MN165577 |
| Onokochanskaya_34g | Lake Baikal, Onokochanskaya Bay (55.57 N 109.23 E) (15 Jul 2018) | MN165578 |
| Sludyanka_52c | Lake Slyudyanskoye (55.46 N 109.16 E) (16 Aug 2016) | MN165579 |
| Zavorotnaya_60g | Lake Baikal,Zavorotnyy Cape (54.28 N 108.49 E) (15 Aug 2016) | MN165580 |
| Tankhoy_69g | Lake Baikal, Tankhoy (51.55 N 105.12 E) (28 Sep 2017) | MN165581 |
| Zavorotnaya_61cg | Lake Baikal,Zavorotnyy Cape (54.28 N 108.49 E) (15 Aug 2016) | MN165582 |
| Senaya_11g | Lake Baikal, Sennaya Bay (52.28 N 105.73 E) (4 Oct 2017) | MN165583 |
| Elohin_3c | Lake Baikal, Elokhin Cape (54.53 N 108.65 E) (1 Oct 2017) | MN165584 |
| 16Yarki | Lake Maloye Akuli (55.71 N 109.91 E) (16 Jul 2018) | MN165585 |
| 5Yarki | Lake Maloye Akuli (55.71 N 109.91 E) (16 Jul 2018) | MN165586 |
| 15Yarki | Lake Maloye Akuli (55.71 N 109.91 E) (16 Jul 2018) | MN165587 |
| 13Yarki | Lake Maloye Akuli (55.71 N 109.91 E) (16 Jul 2018) | MN165588 |
| 10Yarki | Lake Maloye Akuli (55.71 N 109.91 E) (16 Jul 2018) | MN165589 |
| 12Yarki | Lake Maloye Akuli (55.71 N 109.91 E) (16 Jul 2018) | MN165590 |
| 3Yarki | Lake Maloye Akuli (55.71 N 109.91 E) (16 Jul 2018) | MN165591 |
| 4Yarki | Lake Maloye Akuli (55.71 N 109.91 E) (16 Jul 2018) | MN165592 |
| 8Yarki | Lake Maloye Akuli (55.71 N 109.91 E) (16 Jul 2018) | MN165593 |
| Frolikha_16g | Lake Baikal, Frolikha Bay (55.51 N 109.77 E) (17 Aug 2016) | MN165594 |
| Frolikha_14c | Lake Baikal, Frolikha Bay (55.51 N 109.77 E) (17 Aug 2016) | MN165595 |
| Elohin_1c | Lake Baikal, Elokhin Cape (54.53 N 108.65 E) (1 Oct 2017) | MN165596 |
| B_Goloustnoye_17g | Lake Baikal, Bolshoye Goloustnoye (52.02 N 105.40 E) (4 Oct 2017) | MN165597 |
| Elohin_33g | Lake Baikal, Elokhin Cape (54.53 N 108.65 E) (1 Oct 2017) | MN165598 |
| Onguren_28c | Lake Baikal (53.61 N 107.61 E) (3 Oct 2017) | MN165599 |
| Onguren_30c | Lake Baikal (53.61 N 107.61 E) (3 Oct 2017) | MN165600 |
| Onguren_29g | Lake Baikal (53.61 N 107.61 E) (3 Oct 2017) | MN165601 |
| Sludyanka_50c | Lake Slyudyanskoye (55.46 N 109.16 E) (16 Aug 2016) | MN165602 |

| | | |
|---|---|---|
| Kotelnikoskiy_55c | Lake Baikal, Kotelnikovskiy Cape (55.06 N 109.11 E) (16 Aug 2016) | MN165603 |
| Sludyanka_53c | Lake Slyudyanskoye (55.46 N 109.16 E) (16 Aug 2016) | MN165604 |
| Listv_6l | Lake Baikal, Listvyanka (51.85 N 104.86 E) (10 Aug 2018) | MN165605 |
| Zavorotnaya_59c | Lake Baikal,Zavorotnyy Cape (54.28 N 108.49 E) (15 Aug 2016) | MN165606 |
| Sludyanka_51c | Lake Slyudyanskoye (55.46 N 109.16 E) (16 Aug 2016) | MN165607 |
| Sennaya_12c | Lake Baikal, Sennaya Bay (52.28 N 105.73 E) (4 Oct 2017) | MN165608 |
| B_Goloustnoye_19c | Lake Baikal, Bolshoye Goloustnoye (52.02N 105.40 E) (4 Oct 2017) | MN165609 |
| Buguldeyka_23c | Lake Baikal, Buguldeyka (52.52 N 106.05 E) (4 Oct 2017) | MN165610 |
| Yarki_11 | Lake Maloye Akuli (55.71 N 109.91 E) (16 Jul 2018) | MN165611 |
| Yarki_12 | Lake Maloye Akuli (55.71 N 109.91 E) (16 Jul 2018) | MN165612 |
| babushkin_43c | Lake Baikal, Babushkin (51.72 N 105.85 E) (29 Sep 2017) | MN165613 |
| Sennaya_12g | Lake Baikal, Sennaya Bay (52.28 N 105.73 E) (4 Oct 2017) | MN165614 |
| Frolikha_16c | Lake Baikal, Frolikha Bay (55.51 N 109.77 E) (17 Aug 2016) | MN165615 |
| Baykalskoye_71g | Lake Baikal, Baykalskoye village (55.35 N 109.20 E) (1 Oct 2017) | MN165616 |
| Frolikha_15c | Lake Baikal, Frolikha Bay (55.51 N 109.77 E) (17 Aug 2016) | MN165617 |
| Zavorotnaya_60c | Lake Baikal,Zavorotnyy Cape (54.28 N 108.49 E) (15 Aug 2016) | MN165618 |
| Sennaya_13c | Lake Baikal, Sennaya Bay (52.28 N 105.73 E) (4 Oct 2017) | MN165619 |
| Sennaya_13g | Lake Baikal, Sennaya Bay (52.28 N 105.73 E) (4 Oct 2017) | MN165620 |
| Tyya_50 | Tyya River (55.61 N 109.34 E) (16 Aug 2016) | MN165621 |
| Angara_51 | Angara River (52.25 N 104. 28 E) (10 Aug 2018) | MN165622 |
| Angara_52 | Angara River (52.25 N 104. 28 E) (10 Aug 2018) | MN165623 |
| Solzan_4c | Lake Baikal (51.50 N 104.24 E) (28 Oct 2017) | MN165624 |
| Solzan_5c | Lake Baikal (51.50 N 104.24 E) (28 Oct 2017) | MN165625 |
| n-v_r_Osinovka_65 | Lake Baikal (51.50 N 104.24 E) (28 Oct 2017) | MN165626 |
| n-v_r_Osinovka_63g | Lake Baikal (51.50 N 104.24 E) (28 Oct 2017) | MN165627 |

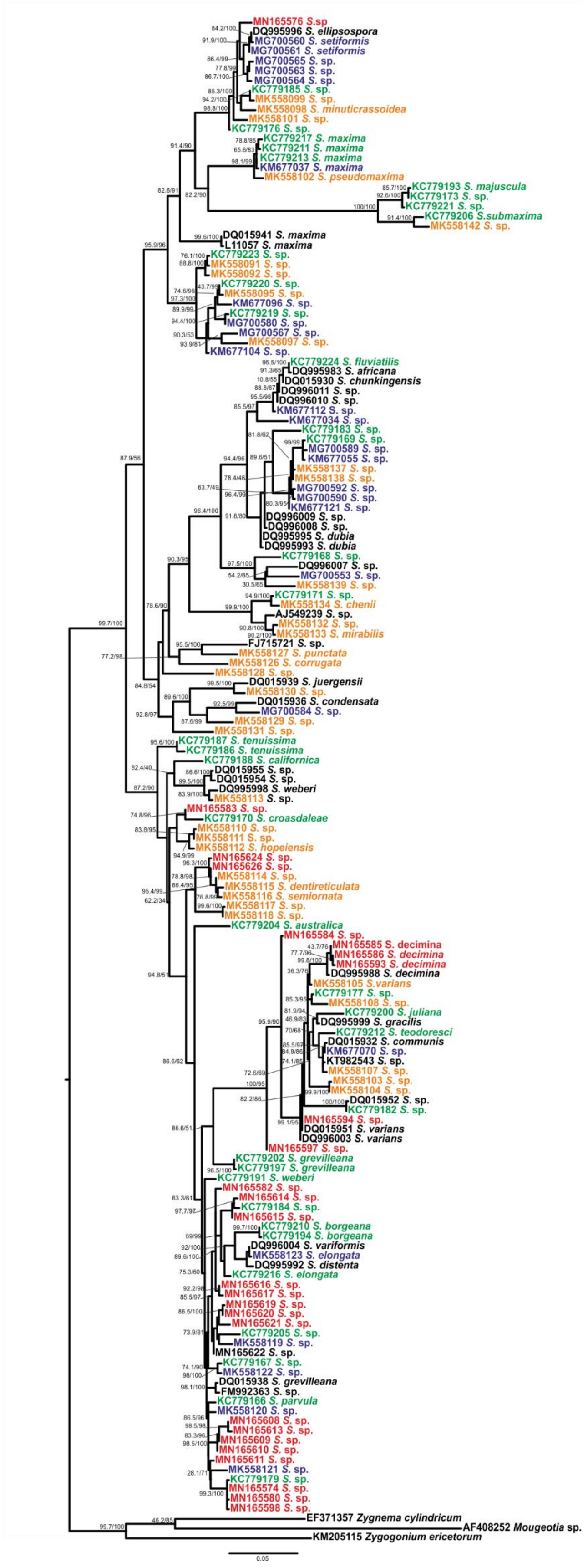


**Figure S1.** Maximum likelihood phylogenetic tree, inferred from the combined DNA data set of *rbc*L containing merged GenBank *Spirogyra* sequences and sequences obtained from single *Spirogyra* filaments in course of the current study. Values at the internal nodes are the alrt and bb supports. Geographic origin of the samples is shown in color: red: Lake Baikal, green: California, blue: Hawaii, orange: Japan, black: unknown

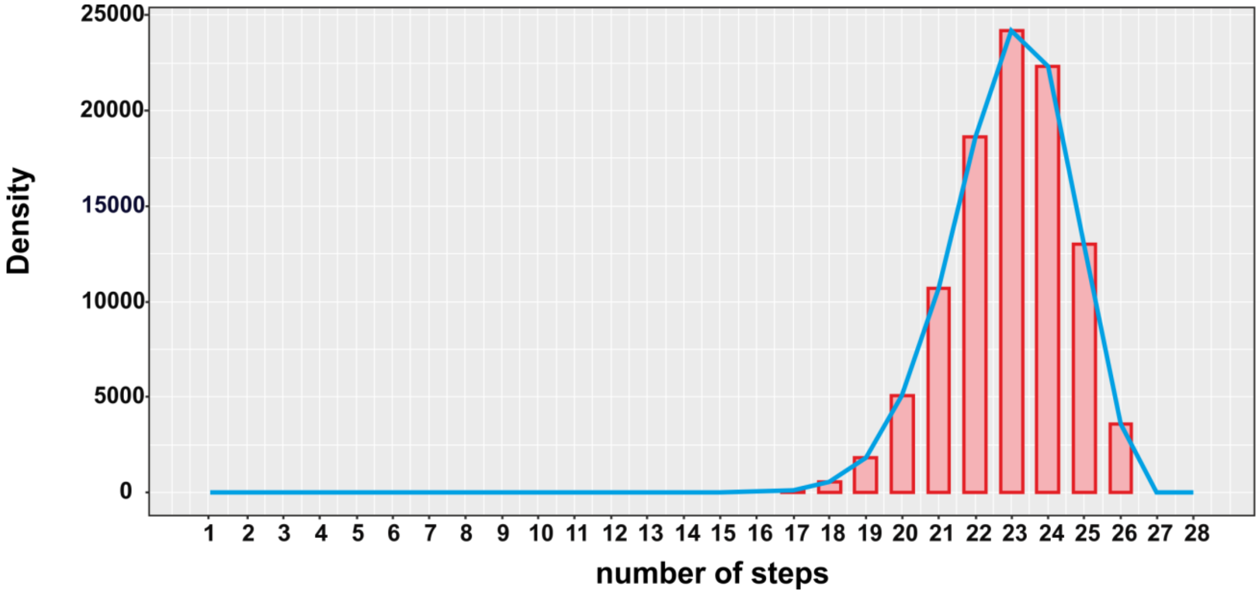


**Figure S2.** Distribution of the minimal number of parsimonious steps after 106 150-steps long random branch-and-bound distortions on the original ML phylogeny. Asterisk indicates the estimated number of steps.
